# Enabling rapid cloud-based analysis of thousands of human genomes via Butler

**DOI:** 10.1101/185736

**Authors:** Sergei Yakneen, Sebastian M. Waszak, Michael Gertz, Jan O. Korbel

**Affiliations:** European Molecular Biology Laboratory (EMBL), Genome Biology Unit, Heidelberg, Germany; EMBL, European Bioinformatics Institute (EMBL-EBI), Hinxton, UK; Institute of Computer Science, Heidelberg University, Heidelberg, Germany.

## Abstract

We present Butler, a computational framework developed in the context of the international Pan-cancer Analysis of Whole Genomes (PCAWG)^1^ project to overcome the challenges of orchestrating analyses of thousands of human genomes on the cloud. Butler operates equally well on public and academic clouds. This highly flexible framework facilitates management of virtual cloud infrastructure, software configuration, genomics workflow development, and provides unique capabilities in workflow execution management. By comprehensively collecting and analysing metrics and logs, performing anomaly detection as well as notification and cluster self-healing, Butler enables large-scale analytical processing of human genomes with 43% increased throughput compared to prior setups. Butler was key for delivering the germline genetic variant call-sets in 2,834 cancer genomes analysed by PCAWG^1^.

---

The cornerstones of Butler design are: full support of multiple cloud environments including academic clouds, declarative infrastructure configuration management, scalable, fault-tolerant operation, and comprehensive monitoring of the data processing. One design principle setting Butler apart from other scientific workflow managers^2-4^ is its heavy reliance on integration of generic open source frameworks to deliver its capabilities. An integration layer brings these together to form a cohesive and comprehensive cloud-computing framework with a focus on genomics applications including in the context of sensitive patient data.

Butler can invoke a variety of analysis algorithms on any cloud. These can either be pre-installed or run as Docker^5^ images, or Common Workflow Language (CWL)^6^ tools and workflows. Each workflow admits parametrisation via JavaScript Object Notation (JSON) configuration files, stored in a database for provenance tracking. Workflow tasks scheduled for execution are deposited into a distributed task queue where available workers can pick them up. This method of operation can scale analyses to thousands of compute nodes.

A main lesson learned from PCAWG^7^ is that genomic data of heterogeneous quality, generated in sequencing centers with varying standard operating procedures and distributed across the globe, frequently suffers from artifacts that lead to numerous compute job failures. These include anomalies due to unusual sequencing library artifacts, failures of bioinformatics tools due to software defects, and inabilities to work with “edge cases” involving sample contamination or non-uniform sequencing coverage, leading cloud-based Virtual Machines (VMs) to fail operating at scale^8^. Delays in recognizing and resolving these failures inevitably reduce the data processing rate, and drive up project duration and costs. Butler enables the timely discovery and resolution of these “expected failures” by providing an operational management toolkit (Supplementary Figure 8) which functions at two levels of granularity – host-level, and application-level. Host-level operational management is facilitated via a health metrics system that collects system measurements at regular intervals from all deployed VMs. These metrics are aggregated and stored in a time-series database within Butler’s central Monitoring Server. A set of graphical dashboards reports system health to users continuously while supporting advanced querying capabilities for in-depth troubleshooting. Application level monitoring is facilitated via systematic log collection and extraction, and logs are then stored in a queryable search index^9^. These tools provide multidimensional visibility into operational bottlenecks and error conditions as they occur, in a manner that is aggregated across hundreds of VMs. A rule-based anomaly detection engine defines normal operating conditions which, when breached, trigger handling routines that can notify the user by sending email, Slack or Telegram messages, and enables automated restarting of offending workflows or entire VMs allowing the cluster to self-heal. These monitoring and operational management capabilities, which prepare cloud users for ‘expected failures’, set Butler apart from current scientific workflow frameworks^6,10-12^(Supplementary Table 3). These capabilities enable highly efficient use of Butler in studies comprising thousands of patient genomes, such as PCAWG, where anomalies and infrastructure failures must be expected, and in fact occur at high rates. These capabilities further facilitate deployment on preemptable infrastructure (such as AWS spot-instance fleets) operating on short-lived VMs that can be shut down by the provider at any time, which are available to users at 10% of the regular cost.

Large genomics data sets are increasingly provisioned on different cloud computing platforms utilizing a mix of public, private, and hybrid clouds^13^. In order to be successful, workflow systems employed in global projects must be flexible in their ability to operate on different environments, including academic clouds, to allow researchers to bring their computational pipelines to the data – in particular, in cases where the raw data itself cannot be moved. The recently developed cloud-based scientific workflow frameworks Nextflow^10^, Toil^11^, and GenomeVIP^12^ focus their support largely on individual commercial cloud computing environments – mostly Amazon Web Services (AWS) – providing incomplete functionality for other major providers (especially academic clouds), which would limit (or prevent) their use in globally pursued studies such as PCAWG that require multi-cloud operation including a major academic cloud component due to practical and regulatory requirements^14,15^. Butler provides full support for operation on OpenStack-based commercial and academic clouds, AWS, Microsoft Azure, and Google Compute Platform, thus enabling the next generation of global disease genomics studies based on hundreds of thousands of patient genomes requiring globally distributed cloud-based data processing in distinct institutions and jurisdictions^14,15^.

We assessed Butler’s ability to facilitate large-scale analyses of patient genomes in the context of PCAWG, where Butler was deployed on 1,500 CPU cores, 5.5 TB of RAM, and 1 PB of storage. We implemented and successfully tested workflows based on BWA^16^, freebayes^17^, Delly^18^, as well as several tools for somatic variant calling including Pindel^19^ and BRASS^20^ using Butler. Utilizing Butler’s suite of workflows we carried out variant discovery and joint genotyping of >90 million germline genetic variants (SNPs, Indels, and SVs) across a 725 TB dataset comprising the PCAWG cohort of 2834 high-coverage tumour as well as normal whole genome sequences. Additionally, we carried out sequence alignment and germline as well as somatic variant calling on 232 high-coverage prostate cancer tumour-normal sample pairs collected within the International Cancer Genome Consortium (ICGC). To accomplish these analyses we executed over 2.5 million compute jobs utilizing 546,552 CPU hours. The management overhead of employing Butler for these analyses was less than 2% of the overall computational cost.

We contrast the observed performance of Butler against the performance of the “core somatic PCAWG pipelines” (Figure 2), which represent the current state-of-the-art in the field achieving almost complete feature parity with other currently available cloud-based scientific workflow frameworks^6,10-12^ (Supplementary Table 3). These PCAWG pipelines used the same IT infrastructure and computed over the same samples – but did not utilize Butler. Our metric to estimate the highest achievable processing rate for an analysis is defined as the smallest proportion of time required to process 5% of all samples, calling it the “target processing rate”. We calculate the ratio of the actual processing rate to the target processing rate (Figure 2 a-b). Butler-operated pipelines were closer to the target processing rate (mean actual/target rate ratio 0.696) than the core PCAWG pipelines (mean actual/target rate ratio 0.490) (Figure 2 c). Butler-based analyses showed a duration 1.43 times the ideal target duration, while core PCAWG pipelines showed a duration of 2.04 times the ideal target duration – which is 43% slower. Additionally, core PCAWG pipelines exhibited a highly non-uniform processing rate (Figure 2 d) deviating 23.1% on average (min 0.0%, max 57.8%, sd 15.0%) from the ideally uniform trajectory of processing 1% of samples in 1% of analysis time, while Butler-based pipelines (Figure 2 e) performed in a significantly more uniform manner only deviating 4.0% (min 0.0%, max 15.6%, sd 3.7%) over the same sample set on average (see Methods). These abilities have resulted in the adoption of Butler for analyses carried out by the international Pan Prostate Cancer Group, and in the European Open Science Cloud pilots (https://eoscpilot.eu).

We have developed Butler to deal with the challenges of working with diverse cloud computing environments in the context of large-scale patient genomic data analyses. The operational management tools provided with Butler help overcome the key challenge that impacts analysis duration – the ability to autonomously detect, diagnose and address issues in a timely manner – thus allowing researchers to spend less time focusing on resolving error conditions and considerably reduce analysis duration and cost. The comprehensive nature of the Butler toolkit sets it apart from current scientific workflow managers^6,10-12^ by offering an efficient and scalable solution for modern global cloud-based big data analyses (see Supplementary Table 3 for feature comparison).

## Methods

Overall, the Butler system is composed of four distinct sub-systems:

### Cluster Lifecycle Management

This sub-system deals with the task of creating and tearing down clusters on various clouds, including defining Virtual Machines (VMs), storage devices, network topology, and network security rules. To fulfill these requirements in a cloud agnostic manner Butler utilizes an open-source framework called Terraform, developed by Hashicorp. Terraform uses a proprietary human-and machine-readable file format for specifying cluster configurations that is called HashiCorp Configuration Language (HCL). Using this language the end user can define a number of constructs for cluster management. The key task of Terraform is to perform Create, Read, Update, and Delete on cluster resources. Running Terraform causes the tool to inspect the current state and compare it to the target state, issuing any necessary commands to update current state to the target. Butler comes with a set of Terraform configuration files that define templates for all of the VMs that constitute a functional Butler cluster, as well as configurations for network security. A typical Butler cluster consists of Control VMs and Worker VMs and templates for both are available. The users are expected to adapt the templates as needed for their use case, providing their own credentials, cluster size, and other configurations.

### Cluster Configuration Management

This sub-system deals with configuration and software installation of all VMs in the cluster. VMs typically will have hundreds of programs installed and configured on them, oftentimes with intricate interdependencies and inter-machine communication requirements. The Saltstack open-source Configuration Management system integrated in Butler allows managing these dependencies and installation details independently of the Operating System (OS) chosen for the Virtual Machines for deployments involving hundreds of servers (Figure 1). The Configuration Management System is controlled by a Master node that acts as the authority on the state of a cluster of Minion nodes. The Master has a set of configuration definitions defined and accessible through a git repository. Each Minion can have a number of roles assigned to it, and the Master maintains mappings between roles and configuration definitions. Once the Master has determined what roles a Minion has it can issue the necessary commands to apply relevant configurations to the Minion. Butler ships with configuration definitions required to run Butler itself as well as those needed to execute the bundled workflows (sequence alignment with bwa, germline variant calling with freebayes and Delly, and somatic variant calling with Sanger Institute’s CGP tools). Additional configurations can be defined by the user as necessary.

**Figure 1:**
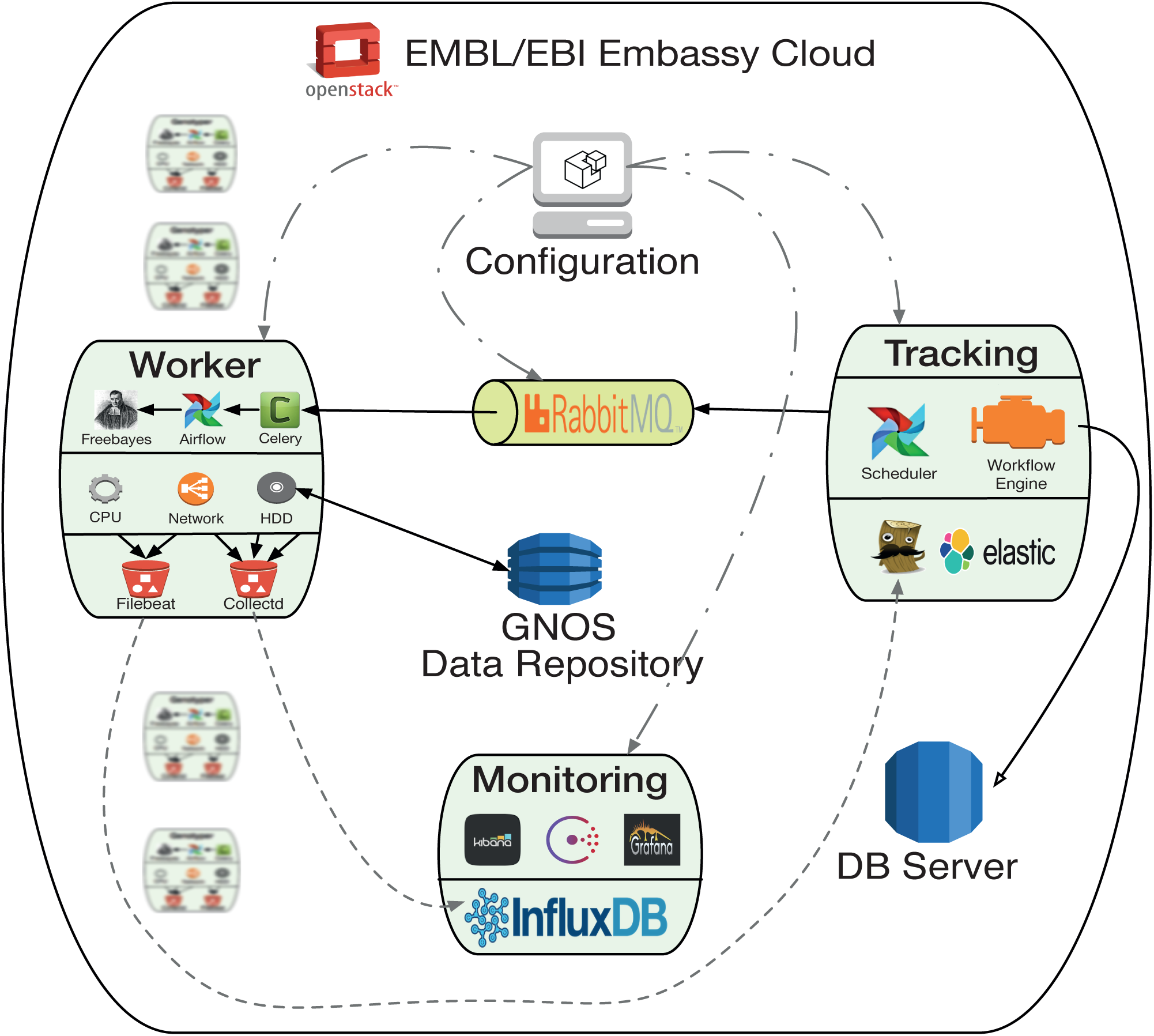
Butler framework architecture as deployed on the EMBL/EBI Embassy Cloud for PCAWG. The Configuration VM manages software installation and settings for all VMs. Tracking VM runs the Workflow Engine and Scheduler which transform workflow definitions into executable tasks and deposit them into a queue. A fleet of Workers run tasks by picking them up from the queue and running the algorithms embedded within the task definition. The Monitoring VM collects health metrics from all other VMs and aggregates them into a set of operational management dashboards.

**Figure 2:**
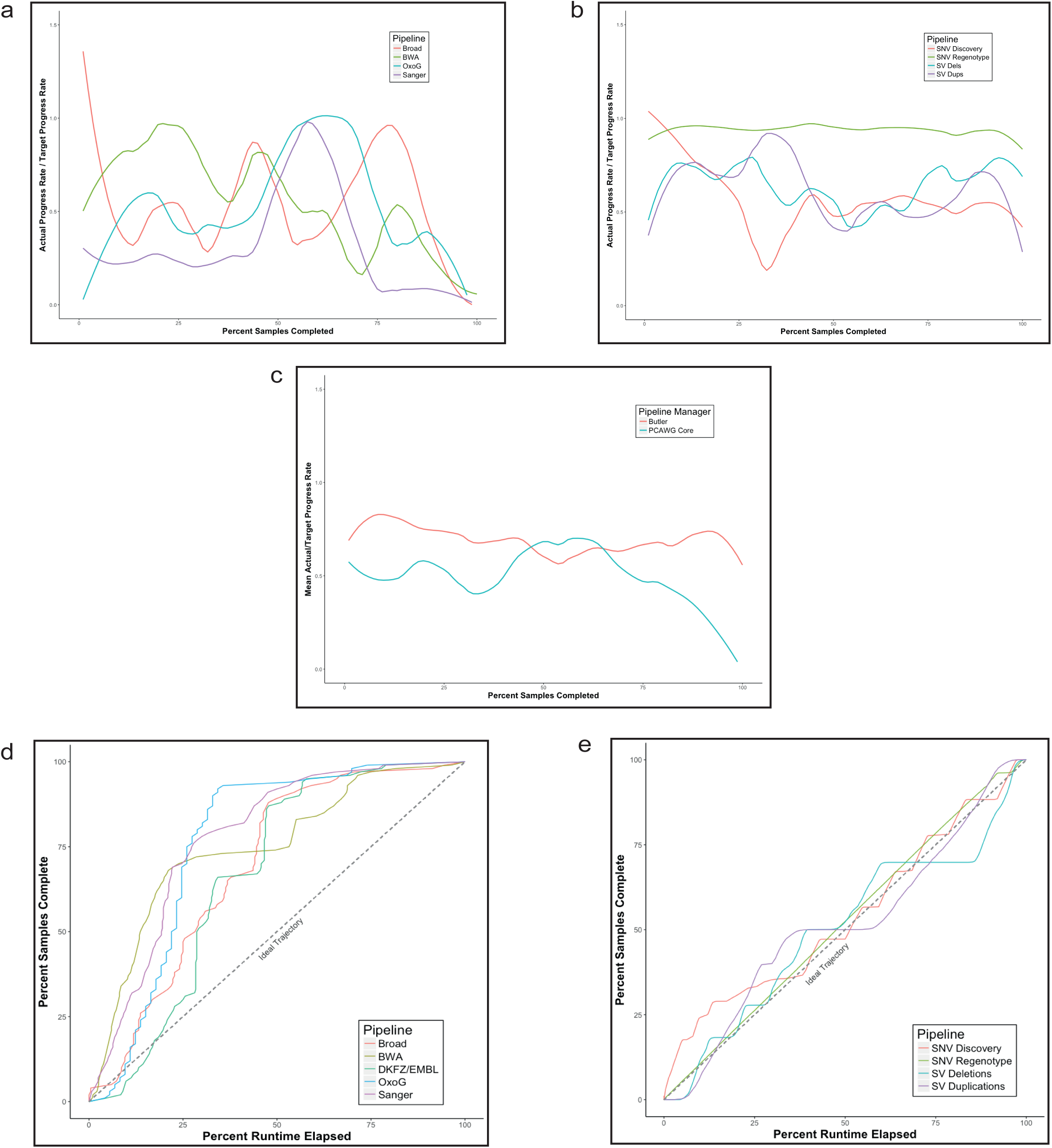
Comparison of analysis progress between “core” PCAWG pipelines and Butler-based pipelines. (a-b) The target progress rate is the minimal proportion of time required to process 5% of samples for each analysis pipeline. We compare the ratio of actual to target progress rates across the cohort for “core” PCAWG Pipelines (a) vs Butler pipelines (b). (c) Mean actual/target progress rate ratio across pipelines for “core” PCAWG (mean 0.49) vs Butler (mean 0.7) pipelines. (d-e) Given a constant allocation of resources the “ideal” progress rate is 1% of samples processed for each 1% of total analysis runtime. All “core” PCAWG pipelines deviate significantly from the ideal trajectory (mean deviation 21.3%, sd 15.0%) taking half of the overall analysis time to complete the last 15-30% of the samples. Butler-based pipelines demonstrate a dramatically lower mean deviation from the ideal trajectory (mean 4.0%, sd 3.7%) over the same sample set.

### Workflow System

The Workflow sub-system is responsible for allowing users to define and run scientific workflows on the cloud. Butler integrates the open-source distributed workflow system called Airflow developed by Airbnb for this purpose. The key component at the heart of Airflow is the Airflow Scheduler (Figure 1). The airflow-scheduler is a service that runs perpetually on a VM and examines the state of all running workflows. All workflow tasks that meet the preconditions for being runnable are immediately “scheduled” for execution. In the context of Airflow scheduling means depositing the task into a queue (running on a separate Queue Server VM) from which a Worker VM (Figure 1) can eventually pick it up. The Worker VMs run an airflow-worker service that periodically polls the task queue for available tasks, and when the task is runnable by a particular Worker, that Worker consumes the task message from the queue and assumes execution. In order to keep track of the status of Workers and workflow execution each Worker periodically sends heartbeat messages to the Scheduler to communicate its state. The state is persisted by the Scheduler to a PostgreSQL database, which runs on a DB Server VM (Figure 1).

The user can communicate with and commandeer Airflow via the Airflow CLI, as well as a Web UI. The Web UI is provided via the airflow-flower, and airflow-webserver services, which can run on the same VM as the Scheduler or on a separate VM, depending on system load. Conceptually, an Airflow workflow is a Directed Acyclic Graph whose vertices represent tasks and edges indicate task sequence. In its implementation an Airflow workflow is a Python program that can use any Python language construct or library. This allows the users to create workflows of arbitrary complexity and functionality.

An Analysis Tracker module is built into Butler in order to allow the user to define analyses, specify what workflows are part of these analyses, and track the status and execution of Analysis Runs - instances of running a particular workflow on a particular data sample within the context of an Analysis (Supplementary Figure 1).

When an Analysis Run is first created it is given a Ready status (Supplementary Figure 2), indicating that it is ready to be scheduled for execution. Once the Scheduler has scheduled the Run for execution it is given a Scheduled status. When workflow execution starts the Run is marked In-Progress. Once the Run is successfully completed it enters a Completed status. If, at any point, the Run encounters an error condition it cannot recover from, the Run Status is set to Error. When the error condition is addressed the Run status should be set to Ready so that it can start from the beginning.

In order to fulfill the workflow configuration and parametrisation requirements Butler implements a tri-level configuration mechanism (Supplementary Figure 3), allowing the user to specify configurations at Workflow, Analysis, and Analysis Run levels. At runtime all three configuration levels are merged into one “effective” configuration that applies within the execution context. Because it is important for configuration to be both human-readable and machine-readable Butler uses the JSON format to encode configuration information. PostgreSQL, in turn, has native support for storage and deep querying of JSON values, thus making it an ideal choice for configuration persistence.

### Operational Management

This sub-system provides tools for ensuring continuous successful operation of the cluster, as well as for troubleshooting error conditions. In general, the Operational Management tools fall into two categories, those that collect observations about the state of each component in the system at runtime, and those that aggregate this data and present it to the user in the form of queryable databases and management reports. We delineate two major sources of data that is indicative of system state - System Metrics, and Server Logs. While metrics provide more of a coarse-grained view of the overall health of a particular Virtual Machine, server logs can give much more of a fine-grained view of the underlying system at an application level, and down to individual lines of code that are running at any given time. Butler has dedicated components for the collection and management of these data sets

Each VM runs a metric collection daemon called collectd, which is an open-source package that is able to make periodic measurements of a large number of system metrics and ship them off to a centralized Monitoring Server. The definition for which metrics are collected is specified in a special configuration file. Because we are interested in observing not only the metrics as they are measured in the present, but also the dynamics of how metric values change over time, we need a mechanism for persisting this information. For this purpose the Monitoring Server component of Butler ships with an instance of a database product called InfluxDB, which is an Open Source database system that is optimized for recording time series data.

The metrics collection system is collecting 50 different metrics per host on average, sampled at intervals of 10 seconds. Given a cluster of 200 Virtual Machines the monitoring system collects and stores 86,400,000 data points in a 24 hour time period. This volume of data is quite difficult for the user to comprehend and make use of, and Butler provides visualization tools to enable the display of aggregate statistics based on the monitoring data using a Graphical User Interface (Supplementary Figure 8). The main goal of the visualizations is to give the user an overview of the trends observed within the compute cluster with respect to a set of representative performance metrics, and to alert the user to any conditions that threaten the health of Virtual Machines and the scientific analyses they run.

Because of the potentially extremely high value of the information contained in server logs, we deploy a system of log harvesting and centralized storage that enables the Virtual Machines that are part of Butler to parse the logs that are being generated locally for interesting events, and send those events to a centralized search index which is amenable to efficient querying and visualization. These open-source tools are known as the ELK stack (Elasticsearch, Logstash, Kibana).

Each Virtual Machine in the cluster runs a log shipper - Filebeat. It is responsible for finding, harvesting, and locally aggregating logs. Logstash runs on a separate centralized server and is responsible for parsing the logs forwarded from Filebeat and sending the parsed information on to the Elasticsearch index. Elasticsearch is a general-purpose scalable text indexing and search engine that supports clustering and sharding of data. Just as it is difficult to grasp and analyze performance metrics due to the number of data-points generated, it is as difficult to grasp log messages from a large cluster. We utilize a similar set of visualization tools to the ones we use for metrics, to solve this problem for server logs within Butler. The Kibana dashboarding framework allows us to create graphical dashboards that visualize log events of interest, as well as providing a web-based query interface to the Elasticsearch log messages index (Supplementary Figure 4).

The Monitoring Server runs an anomaly detection and alerting library called Telegraf which defines a series of rules that specify the normal operating conditions for the cluster, such as all hosts regularly responding to ping, CPU below 80%, disk utilization below 85%, workflow tasks sending heartbeats every minute etc. Coupled with the application metrics gatherer statsd, the system builds an empirical distribution of the duration of various workflow tasks and knows when tasks take longer than they historically have. The anomaly detection software monitors the time-series database that records all metrics and periodically evaluates the rule-set against it. When the system detects a breach of one or more rules it can take remedial action, such as sending a warning email, a message to a Slack topic, a Telegram, or schedule the restart of a particular workflow or the reprovisioning of a particular VM. These abilities allow Butler users to be always up to date about the health of the system and allow Butler clusters to self-heal when they encounter problematic scenarios.

The Butler framework consists of many different services that reside on a number of different servers and need to be able to communicate with each other. To accomplish this in a flexible manner we needed to establish a Service Registry so that IP addresses of servers that host particular services could be looked up by service name. To accomplish this Butler uses an open-source service discovery framework called Consul. Consul provides a cross-data-center distributed Service Name Registry that is available via HTTP and DNS protocols. In addition to registry capabilities Consul provides basic health checks for the underlying services, testing whether the IP and port the service is supposed to be listening on are actually reachable.

### Butler Deployment

Butler has been validated for production use on the EMBL-EBI Embassy Cloud - an academic cloud computing center that runs an OpenStack-based environment (Figure 1). The Embassy Cloud plays a key role in the PCAWG project by donating substantial storage and cloud computing capacity over the course of 3 years. The total amount of resources dedicated to the project by the Embassy Cloud is:

- 1 PB Isilon storage shared over NFS
- 1500 compute cores
- 6 TB RAM
- 40 TB local SSD storage
- 10 Gb network

These resources have been used to host one of the six PCAWG data repositories that exist worldwide, as well as performing a number of scientific analyses for the project. We have used Butler extensively on the Embassy Cloud in order to carry out the analyses for the Germline Working Group of PCAWG.

To deploy Butler on the 1500 core cluster we set up five different profiles of VMs, each playing a number of different roles (Supplementary Table 1).

We observe that the mean *r*_*opt*_ is significantly higher for Butler-based pipelines at 0.46 than for the core PCAWG pipelines at 0.13 (Supplementary Table 2). For each pipeline and each 1% of the samples under analysis, we then compute a metric *e* (for effectiveness), defined as the proportion of *r*_*opt*_ actually achieved.

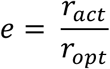

Comparing the core PCAWG and Butler pipelines with respect to *e* (Figure 3 a-c) we observe that effectiveness is on average lower for PCAWG pipelines (*μ*_*e*_*PCWAG*__ = 0.49) than Butler pipelines (*μ*_*e*_*PCWAG*__ = 0.70). Assessing the expected analysis duration for the two sets of pipelines we observe:

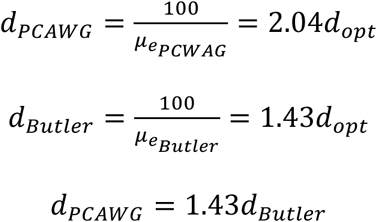

Thus, the expected duration for PCAWG pipelines is 43% longer than that for Butler-based pipelines.

We wish to further compare core PCAWG pipelines with Butler pipelines on the basis of uniformity of rate of progress through an analysis. Given a constant resource allocation an ideal analysis execution processes 1% of all samples in 1% of the analysis runtime. We divide the sample set into 100 equal size bins and measure the percentage of overall analysis time spent on processing each bin (Figure 3 d-e). Deviations from the diagonal indicate inefficiencies in data processing. Measuring this deviation we observe that PCAWG pipelines deviate 23.1% from the diagonal on average (min 0.0%, max 57.8%, sd 15.0%) while Butler pipelines only deviate 4.0% (min 0.0%, max 15.6%, sd 3.7%) from the diagonal on average, over the same sample set. This indicates that Butler pipelines are impacted less by various causes that slow down an analysis (such as job and infrastructure failures).

### Data Availability

All of the data is made available through the PCAWG data portal (https://dcc.icgc.org/pcawg) as described in detail in the PCAWG marker paper^1^.

### Code Availability

The source code for Butler is freely available at http://github.com/llevar/butler under the GPL v3.0 license.

## Acknowledgements

The authors would like to acknowledge the EMBL-EBI Embassy Cloud, The Cancer Genome Collaboratory, Amazon Web Services (AWS), Google Compute Platform, and Microsoft Azure for providing compute infrastructure and cloud resources.

We kindly acknowledge the following EMBL-EBI employees for the invaluable assistance with the EMBL-EBI Embassy Cloud: Andy Cafferkey, Charles Short, David Ocaña, Dario Vianello, Erik van den Bergh, Steven Newhouse, Ewan Birney.

JOK was supported by the European Open Science Cloud Pilot study (European Commission award number: 739563) and the BMBF (de.NBI project: 031A537B). SW was supported through a SNSF Early Postdoc Mobility fellowship (P2ELP3_155365) and an EMBO Long-Term Fellowship (ALTF 755-2014).

## Author Contributions

S.Y. designed, implemented, and executed Butler for the analyses described in this manuscript.

S.M.W. designed workflows, and assessed the integrity of the framework.

S.Y. and S.M.W. performed the data analysis.

The manuscript was written by S.Y. and J.O.K., with input from all authors.

